# *TraitCorr* – correlating gene expression measurements with phenotypic data

**DOI:** 10.1101/557975

**Authors:** Thomas Nussbaumer, Christian Wagner, Parviz Heidari

## Abstract

Today, transcriptomes and microarrays can be generated and analysed in high quantity. In addition, experiments often include descriptive information about each sample which needs to be compared to the gene expression profiles. The understanding of the relationships between gene expression and phenotype is introduced as new challenge in system biology. Combining expression (RNA-seq and microarray) and phenotype data could reveal the role or effects of each gene on traits. To address all these needs, the user-interface *TraitCorr* was developed which allows to determine genes that are significantly correlating with a selected trait. Furthermore, it allows to determine significantly correlated genes among different traits and provides visualisation and analysis possibilities.

## Introduction

Transcriptomic datasets summarize the abundance of thousands of genes in a particular species under a given condition. However, to fully understand an experiment and to detect most relevant genes in the dataset, it is necessary to compare the abundance measurements to phenotype information such as the severity or duration after infection when dealing with host-microbe dual RNA-seq or gene array datasets while in clinical datasets it might be interesting to compare the gene expression to the age or disease severity of a patient. After gene expression measurements are normalized, there is the need to identify genes that cause a specific phenotype, thus showing a strong association between the abundance of a gene or protein and trait. This is needed in studies where data doesn’t qualify the required conditions for a genome-wide association study (GWAS) of having a sufficient amount of samples while in addition might not contain a clear control-disease condition for performing differentially expression analysis. The search for relevant genes is similar to a search for a needle in a haystack - as only a few genes remain after intrinsic filter criteria are applied and following the correction for multiple testing and functional analysis is needed to understand the interplay of these genes.

Among tools that allow to correlate expression information with traits is the Weighted gene co-expression network analysis (WGCNA) [1] where trait files can be additionally provided and can be compared to previously calculated modules of co-expressed genes. These correlations can also be computed and compared in particular quantitative trait loci [2]. Other tools have tried to improve the issue of the multiple testing aspects of the obtained correlations between expression patterns and traits [3].

Also several tools and methods exist that linked gene expression with pathway and phenotype information as shown in Papatheodorou et al. [4], while Wang et al. [5] who determined marker genes using gene ranking methods and Ficklin et al. [6] tried determining gene sets to determine association with phenotypes among many other approaches.

Here, with our tool *TraitSec* we integrate abundance data and allows the comparison with phenotypic data along with using different multiple testing corrections. Thereby, it helps the researcher to easily compute correlations between expression information and phenotypic data by offering different correlation coefficient approaches, multiple testing corrections and visualization that are described below.

## Material and Methods

### Graphical user interface for data integration and analysis

The graphical user-interface of *traitCorr* is created with help of the R Package Shiny [7]. The *traitCorr* contains three main panels to analyse and to integrate input data: the *first panel* allows to integrate the two input data files (normalized gene expression of the entire transcriptome and the phenotype information data about the samples as a second data input). Within the *second panel*, correlations can be computed between the entire gene repertoire and a selected trait. In the *third panel*, various phenotypes on the basis of significantly correlated genes can be compared.

### Computation of significant correlation with trait data

Correlations coefficients are compared using the R function *‘cor.test’* between the abundance data and phenotype information. In this comparison, either *Spearman* or *Pearson* correlation coefficient can be selected - allowing to infer a *p*-value and to compute the correlation coefficient which can be then visualized in a customized plot. This plot shows the *p*-values and correlation coefficient from all genes when compared to the selected trait (Figure 2). For the export of the final report, the package ‘*xlsx’* is used where significantly positive or negative correlation coefficients are summarized and *traitCorr* calculates then significantly correlating genes per traits. Once the calculation is ready, the significant genes are stored in the memory. The genes that significantly correlate with a trait are recalculated if the parameter is changed.

### Overlaps of genes correlated significantly between different phenotypes

Overlaps between the various candidate genes are created with the *venn.diagram* R package [8] by selecting phenotype information which can be compared on the basis of common significantly correlated genes using the function *draw.pairwise.venn* and *draw.triple.venn*. It is possible to visualize two or three traits when only genes are shown, that correlated with <0.05 or <0.0001 after multiple testing correction with either Benjamini-Hochberg (BH) or Bonferroni correction.

### Download functionalities

Genes, that correlate significantly with a trait can be exported within the graphical user interface with the help of the *‘xlsx’* package [9]. They can also be exported as a report by using Officer [10] – an R package that provides an overview of all summarized from the analysis.

## Results and Discussion

The *traitCorr* consists of three main panels: While in the *first* panel, a user can upload the two datasets (gene expression data and a second input file containing the phenotype information), in the *second panel* of the GUI correlations can be computed while in the *third panel* different phenotypes can be compared based on genes that significantly correlate with genes (Figure 1).

**Figure 1.**
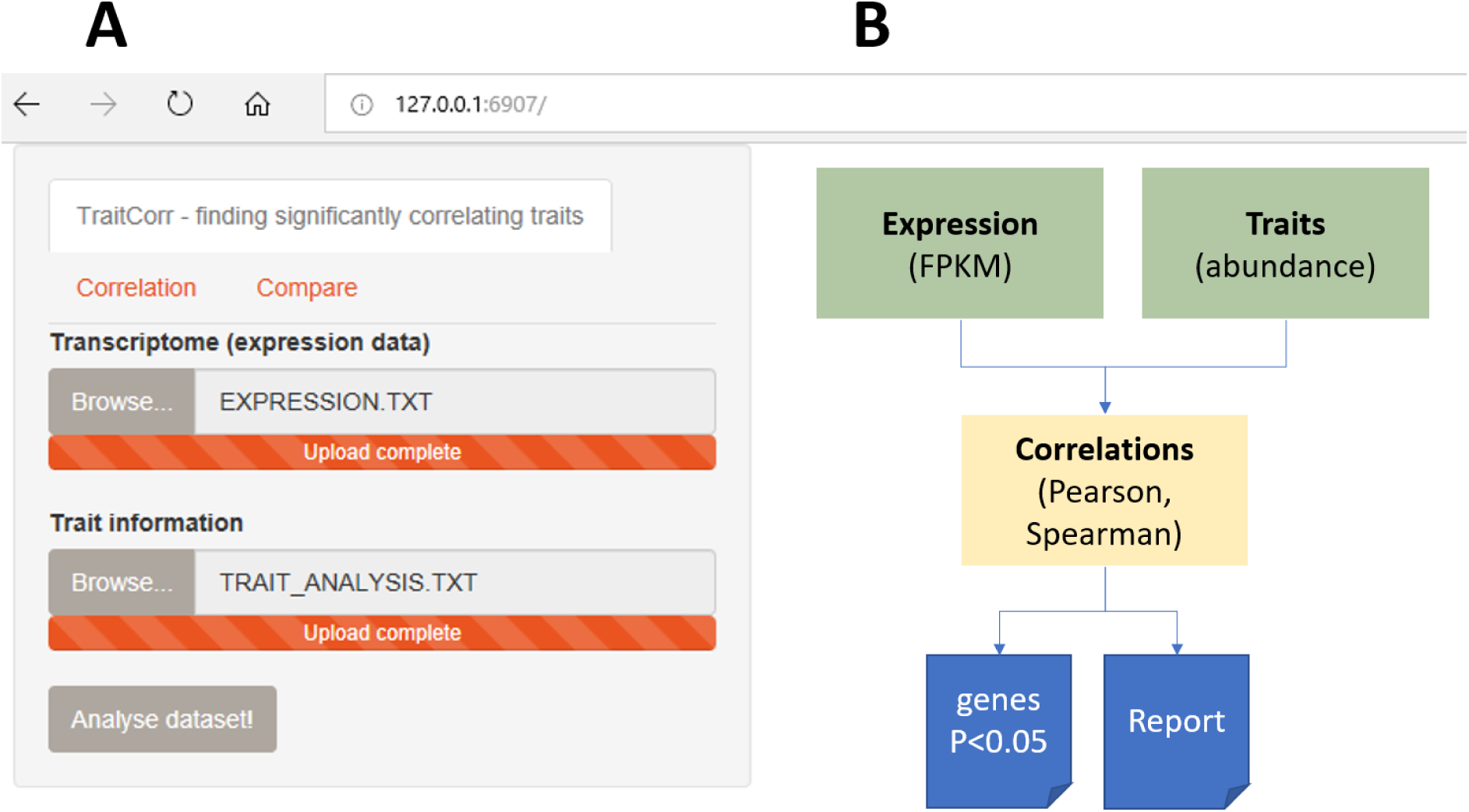
*traitCorr* with three panels to include input, to compute correlations for one particular trait, to compare different phenotypes with each other on the basis of common significantly correlating genes and for downloading significantly correlated genes (subfigure A) and the workflow of *traitCorr* (subfigure B).

If both datasets are supplied, it is possible to calculate correlations between traits and single genes or species by considering either Benjamini-Hochberg (BH), Bonferroni correction for multiple testing correction and by considering *Spearman* or *Pearson* correlations (Figure 2). For all the gene-to-phenotype comparisons, the overall obtained *p*-values and correlations coefficients are shown which also includes the amount of genes that are significantly correlating under a 0.05 level or 0.001 or the genes that are not at all correlated. Plots also depict whether these genes are either positively or negatively correlating including the indications of a strong positive, weak positive, strong negative and weak negative correlation. Furthermore, if different phenotypes are to be compared, the significantly correlated genes are calculated and are visualized as a Venn Diagram to illustrate common genes (Figure 3). Finally, it is possible to extract the genes in each Venn Diagram sector including, if provided in the first step, additional descriptive information about the gene entry.

**Figure 2.**
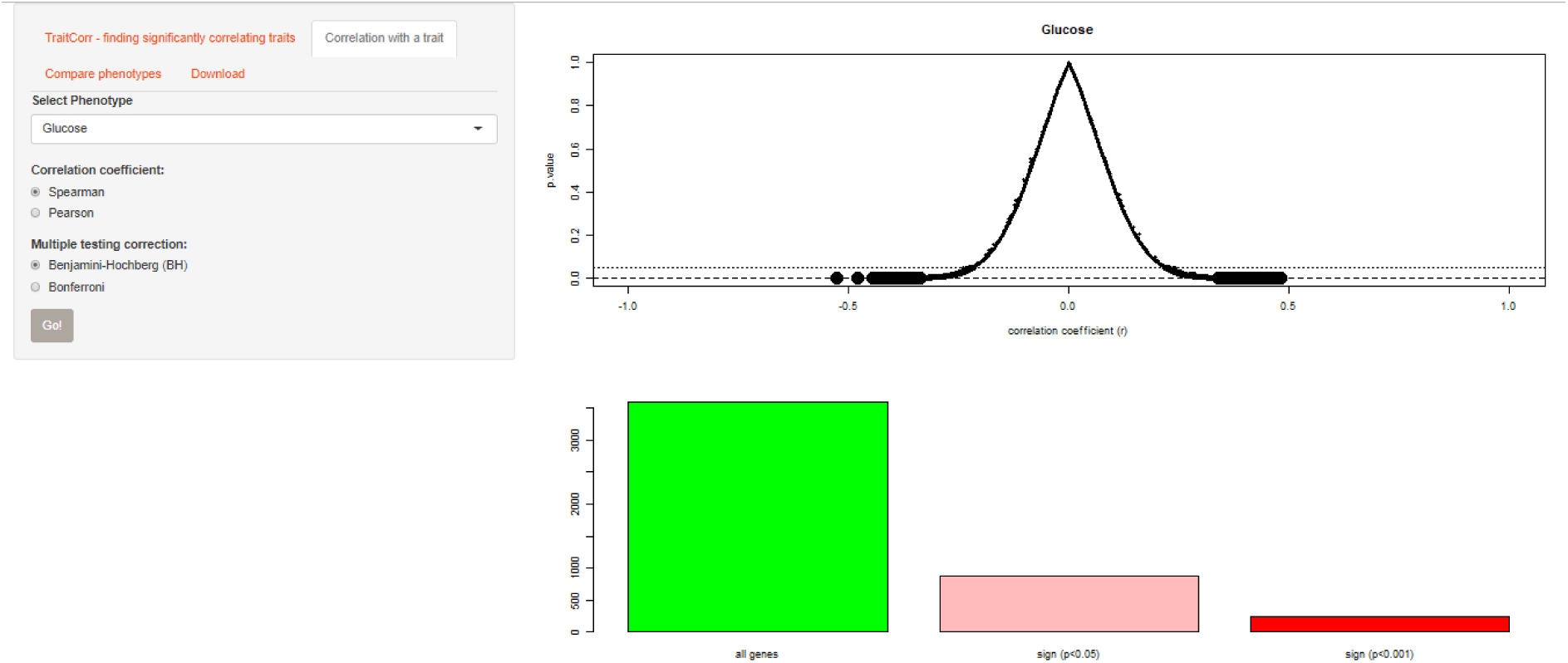
Overview of the correlations coefficients compared to the significance level for each trait. In this example, the trait Glucose was compared to the entire gene repertoire *of the mice gene array data*. The first horizontal lines represent the 0.05 significance level, while the second lower line represents the 0.001 level based on the multiple testing correction after Bonferroni correction.

**Figure 3.**
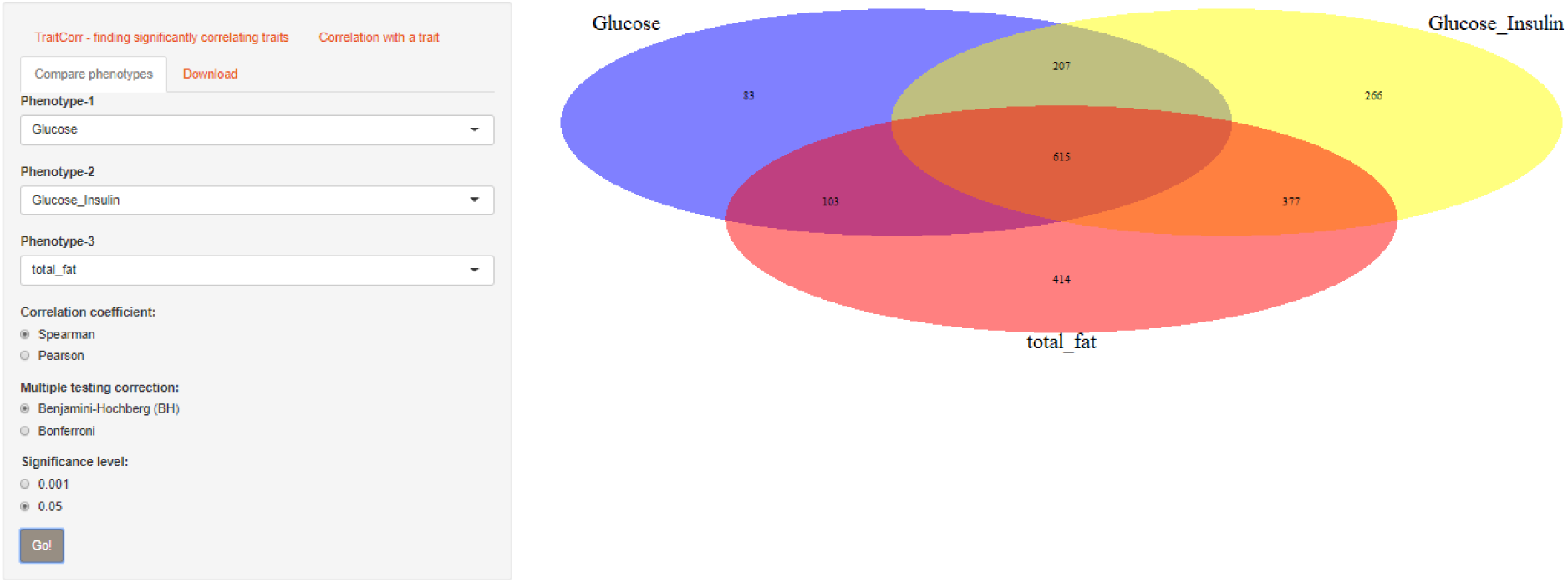
Comparison between the significantly correlating genes for traits Glucose, Glocuse_Insulin and total_fat concentration from Ghazalpour et al. [11].

### *Use case*: mice transcriptome and trait information

In our use case, we have used the demo material that was also used in WGCNA [1] for comparing microarray data of F2 mice with selected traits from Ghazalpour et al. [11]. In our example we have used the data to determine genes that correlate significantly with the abundance of Glucose in Figure 2 where 878 genes are significantly correlated with the trait when using Spearman and after Bonferroni multiple testing adjustment compared 1008 genes with Pearson while we also used the feature “Compare phenotypes” (Figure 3) to compare the significantly correlated genes between the traits “Glucose”, “Glucose_Insulin” and “high_fat” where we found 615 to be shared including 83 genes specific to Glucose. All the data files can be downloaded and further analysed.

## Availability

The R package can be obtained from the following URL: https://github.com/nthomasCUBE/traitCorr.

## Acknowledgments

Dr. Jimmy Omony (PGSB, Helmholtz Zentrum Munich) is acknowledged for critical reading of the manuscript.

